# Functional GPCR expression in eukaryotic LEXSY system

**DOI:** 10.1101/2023.05.04.539202

**Authors:** Aleksandra Luginina, Ivan Maslov, Polina Khorn, Oleksandr Volkov, Andrey Khnykin, Pavel Kuzmichev, Mikhail Shevtsov, Anatoliy Belousov, Dmitrii Dashevskii, Daniil Kornilov, Ekaterina Bestsennaia, Johan Hofkens, Jelle Hendrix, Thomas Gensch, Vadim Cherezov, Valentin Ivanovich, Alexey Mishin, Valentin Borshchevskiy

## Abstract

G protein-coupled receptors (GPCRs) represent an important class of drug targets, and their structural studies facilitate rational drug discovery. However, atomic structures of only about 20% of human GPCRs have been solved to date. Recombinant production of GPCRs for structural studies at a large scale is challenging due to their low expression levels and stability. Here we tested the eukaryotic system LEXSY (*Leishmania tarentolae*) for GPCR production. We expressed the human A_2A_ adenosine receptor (A_2A_AR) in LEXSY, purified it, and compared with the same receptor produced in insect cells, which is the most popular expression system for structural studies of GPCRs. The A_2A_AR purified from both expression systems showed similar purity, stability, ligand-induced conformational changes and structural dynamics, with a remarkably higher protein yield in the case of LEXSY expression.

## Introduction

Structural biology helps to establish the relationship between protein structure and function ^1^, which aids in the rational design of drugs ^2,3^. Among the most important and challenging for structural studies drug targets are membrane proteins, and in particular, G-protein-coupled receptors (GPCRs), the largest membrane protein superfamily in the human genome ^4,5^. GPCRs are complex molecular machines with diverse ligand binding pockets and activation mechanisms that are not yet fully understood ^6^. Due to the involvement of GPCRs in fundamental physiological processes in the human body, they have been implicated in numerous pathologies ^7^. Therefore, understanding the structure and molecular mechanisms of GPCR function is not only of fundamental biological interest but also has enormous potential for improving human health.

Recombinant expression represents one of the bottlenecks in structural and functional studies of GPCRs. Typically, structure determination requires milligrams of properly folded, functional and monodisperse purified protein. Hence, a suitable expression system should be selected for each specific target ^8^. To date, the most popular expression system for GPCR structure determination has been the insect-based baculovirus system (BV) that is responsible for 85% of all recombinant GPCR structures (Fig. 1, Supplementary Table 1). The most common insect cells for BV expression are *Spodoptera frugiperda* (Sf9, Sf21 strains) and *Trichoplusia ni* (Hi5 strain) ^8^. The second most popular system is mammalian cell expression accounting for 13% of recombinant GPCR structures.

**Figure 1.**
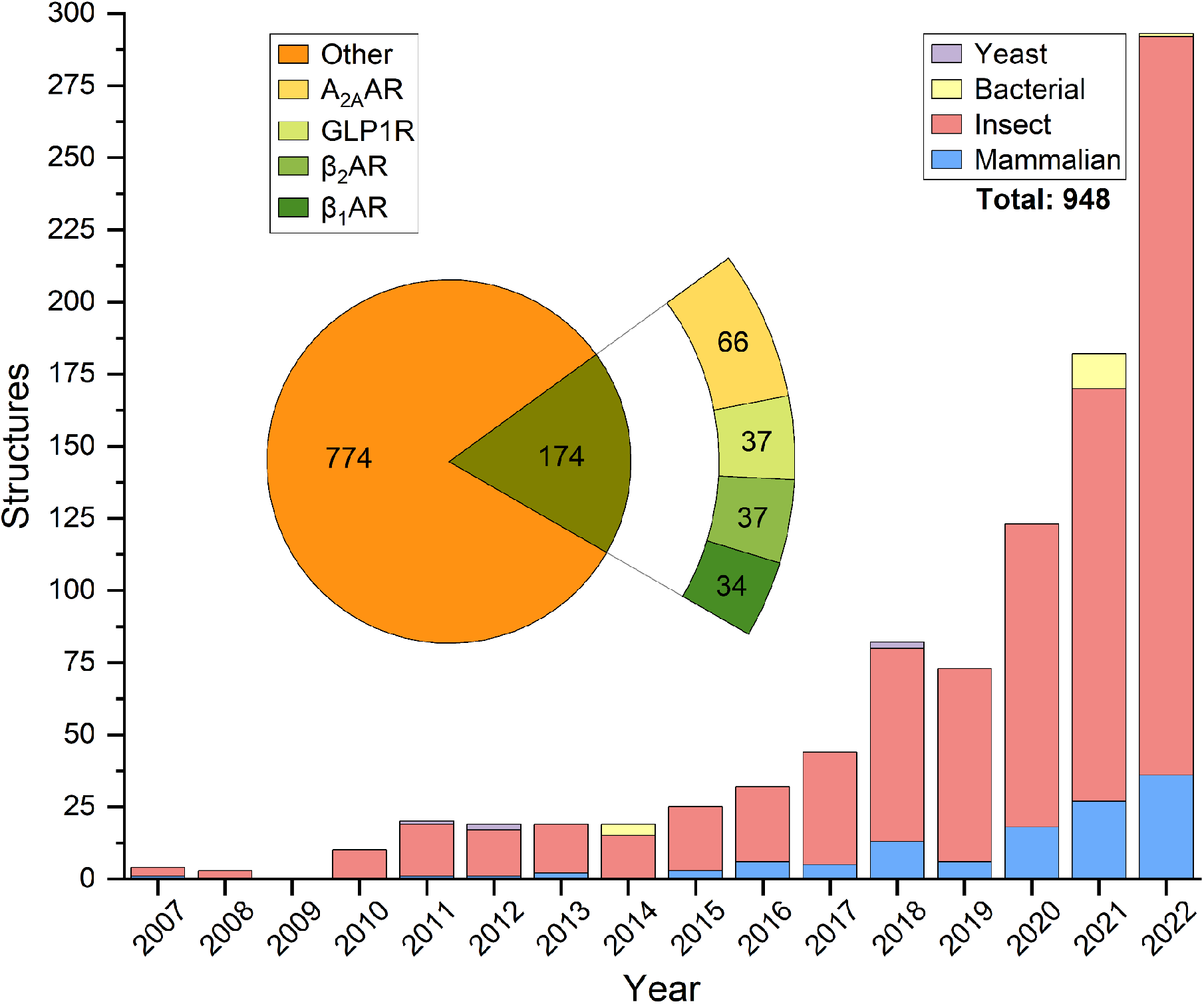
Available structures of receptors by release year and heterologous expression system. Inset: a pie chart showing the distribution of 3D structures for the four most common receptors: A_2A_AR, glucagon-like peptide 1 receptor (GLP1R), β2 adrenergic receptor (β_2_AR), and β1 adrenergic receptor (β_1_AR). Based on GPCRdb^57^ data with correlation to RCSB PDB ^58,59^ for expression system information*. Accessed on March 9, 2023. *In some rare cases, PDB ID-related articles were manually analyzed to determine the expression system.

In this work we evaluated LEXSY as an alternative expression system for GPCRs. LEXSY is based on *Leishmania tarentolae*, a trypanosomatid protozoan nonpathogenic blood parasite of the white-spotted wall geckos. Structures of two transmembrane proteins, channelrhodopsin 2 (ChR2) from *Chlamydomonas reinhardtii* ^9^ and light-driven proton pump LR (Mac) from *Leptosphaeria maculans* ^10^, were recently determined using the LEXSY expression system ^10^. Both of them are retinal 7TM proteins structurally similar to GPCRs, which inspired us to apply this expression system for GPCRs ^11^. However, no systematic validation of LEXSY for GPCR expression has been done so far.

We chose A_2A_AR to benchmark LEXSY for GPCR expression and purification. A_2A_AR is one of the most studied members of the GPCR family. A_2A_AR structures have been captured in inactive, partially active, and fully active states ^12–16^. Three different expression systems: yeast ^17,18^, insect ^19^, and mammalian ^20^ — the most of any GPCR — have been used for A_2A_AR structure determination. Overall, A_2A_AR contributes to around 10% of all recombinant GPCR structures (Fig. 1). Additionally, A_2A_AR has been a workhorse for establishing new biophysical methods for structural studies of GPCRs ^18,21–25^ and is often used as a positive control for purification of other receptors ^26^. Such popularity can be explained by its relatively high stability and yield: over a milligram per liter using BV in the Sf9 cell line.

Here we compared LEXSY with the traditional BV expression system in Sf9 cells for expression, purification, and characterization of the human A_2A_AR. The purified protein samples were characterized by gel electrophoresis (SDS-PAGE), analytical size-exclusion chromatography (SEC), and thermal stability assay (TSA). We further confirmed via single-molecule Förster Resonance Energy Transfer (smFRET) that the extracellular ligand binding translates into identical conformational changes and structural dynamics of the intracellular side of the receptor regardless of the expression system used.

## Results

We expressed A_2A_AR in Sf9 and LEXSY followed by protein purification and functional characterization. In both cases we obtained a monodisperse and functional protein: 1 and 3.6 mg of purified receptor per liter of cell culture of Sf9 and LEXSY, respectively. Protocols for expression and purification were adapted from published structural studies of A_2A_AR for the receptor produced in insect cells ^19^ and of microbial rhodopsins ^9,10^ for the receptor produced in LEXSY. The general applied pipelines are shown side-by-side in Figure 2 and described below.

**Figure 2.**
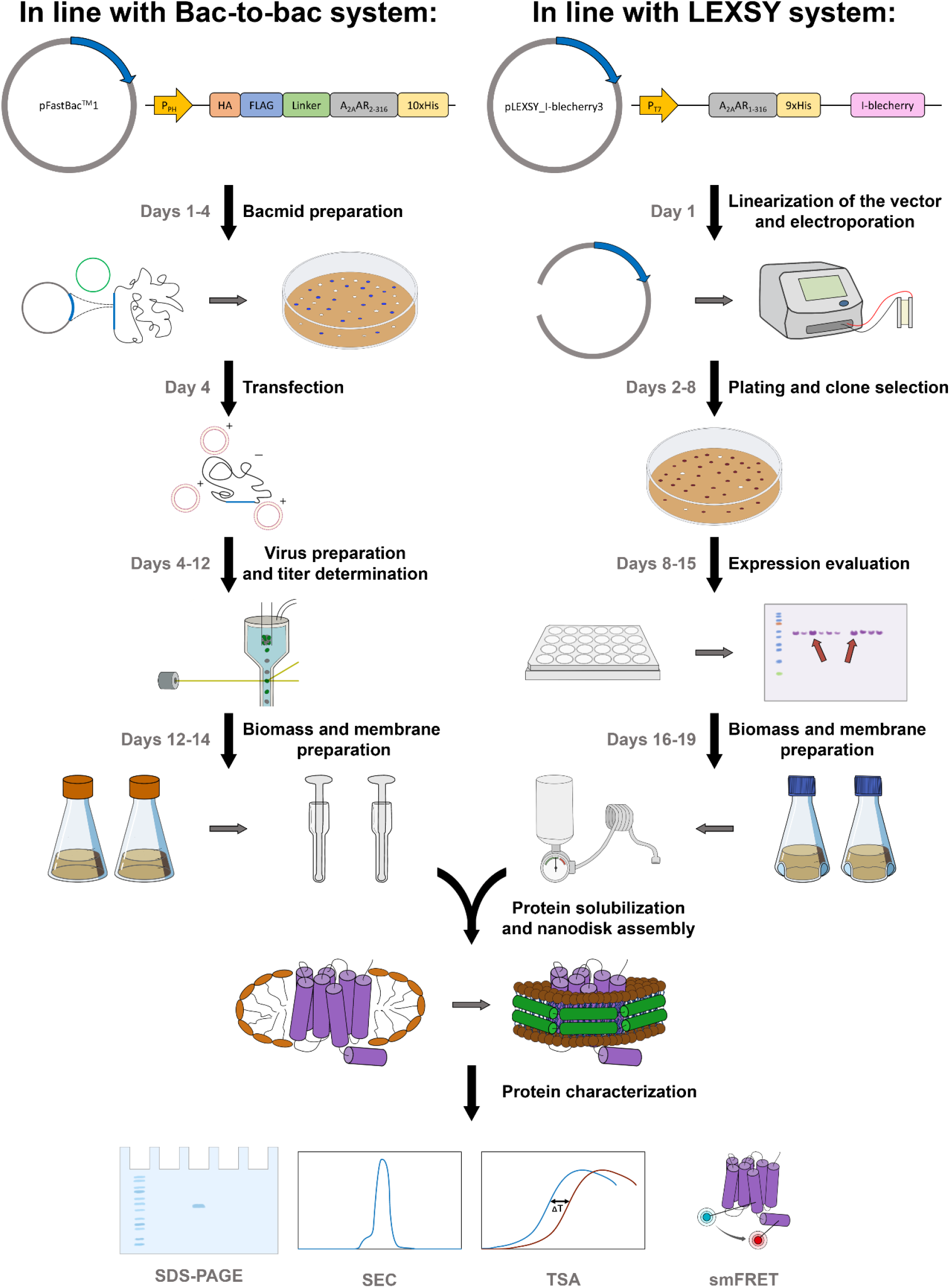
Comparison of production A_2A_AR from Sf9 and LEXSY expression systems. The figure presents a pipeline of the main work stages, including construct design, protein expression, purification, and functional characterization, along with the corresponding timeline. Additional details for each stage are provided in the Results section.

### Construct design and expression vectors

In the case of Sf9, we modified the expression cassette to include an N-terminal haemagglutinin signal peptide (KTIIALSYIFCLVFA) for the cell surface delivery, a FLAG-tag epitope (DYKDDDDK) for measuring surface expression by flow-cytometry, a linker (AMGQPVGAP), a gene of A_2A_AR with a truncated C-terminus (residues 2-316) ^27^, and a C-terminal 10-His-tag. The cassette was cloned into a pFastBac1 (Invitrogen, USA) vector.

In the case of LEXSY, the A_2A_AR gene (1-316) was followed by a GT linker (*KpnI* restriction site used for cloning) and a C-terminal 9-His-tag. The protein gene was cloned into a pLEXSY_I-blecherry3 plasmid. The snake-plots for both constructs are shown in Figure 3.

**Figure 3.**
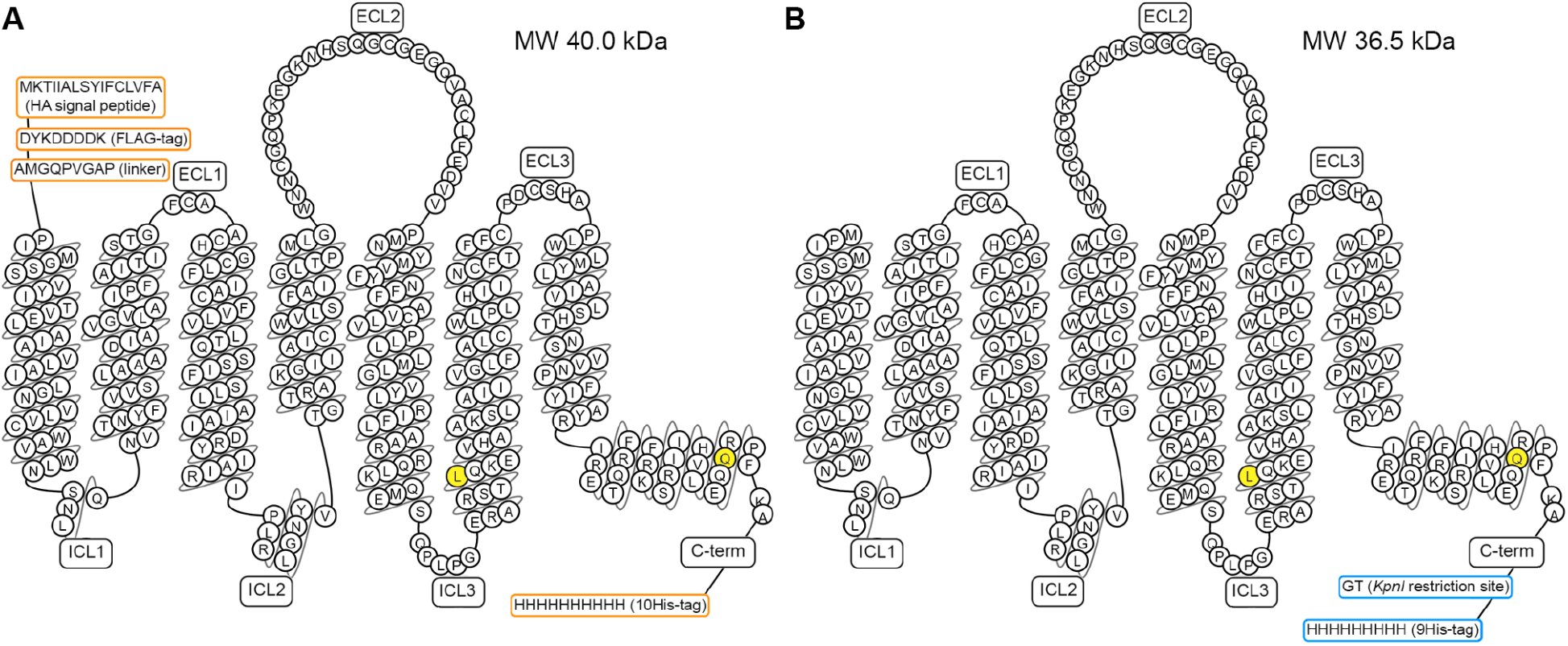
Snake-plot presentation of protein constructs of A_*2A*_AR, expressed in Sf9 (A), and LEXSY (B). L225^6.27^ and Q310^8.65^ residues mutated to cysteines for the labeling for smFRET experiments are colored in yellow. The snake-plot was drawn using the GPCRdb.org website ^57^.

### Protein expression

In the case of Sf9, we transferred the expression cassette into a bacmid upon the process of site-specific recombination in supplementary *E*.*coli* DH10Bac strain cell line. We further transfected Sf9 cells with the bacmid using a cationic lipid reagent and generated baculovirus particles according to the Bac-to-Bac system protocol. To obtain a viral stock with a titer of active virions of >10^9^ particles per mL, we infected the intact Sf9 cell culture. The A_2A_AR surface expression measured by flow-cytometry was 64% with the total expression of 81% (Supplementary Fig. 1A-B).

The overall process from the plasmid DNA to biomass production takes about two weeks. Days 1-4 bacmid preparation: day 1 -DH10Bac transformation; day 3 - transformants inoculation into liquid medium; day 4 - bacmid purification. Day 4-12 virus preparation: day 4 - transfection; day 8 - P0 virus collection and P1 virus infection; day 11 - P1 virus collection and virus titer determination initiation, day 12 - virus titer determination. Day 12-14 biomass preparation: day 12 - biomass infection; day 14 - biomass collection. Recombinant baculoviruses can be stored at +4 °C for several months. Before their subsequent use, the viral stock should be retitered, and if the virus is still viable, expression can be done within two days.

In the case of LEXSY, *Leishmania tarentolae* cells of the inducible T7-TR strain were transfected with the A_2A_AR expression plasmid linearized by the SwaI restriction enzyme. After the clonal selection by western blotting, the transfected cells were grown at 26°C in the dark in media supplemented with Hemin. When OD_600_ = 1 was reached, protein expression was induced by addition of tetracycline, and incubation continued for an additional 24 h, after which cells were collected by centrifugation.

The overall process from plasmid DNA to biomass production takes about two to three weeks. Day 1 - cell transfection (vector linearization and electroporation). Day 2-8 plating and clone selection: day 2 - transfectant plating; day 6 - induction on plates; day 8 - selection of clones. Day 8-15 expression evaluation: selected clones transfer to 24 wells plates; day 11 - induction of clones in 24 wells plates; day 15 - expression control with western blotting (Supplementary Fig. 1C), preculture preparation. Day 16-19 biomass preparation: day 16 - large-scale inoculation of preculture, day 17 - large-scale induction, day 19 - biomass collection. Successfully transformed clones can be frozen. Reestablishing the cell line after thawing and subsequent biomass production takes 15-19 days, including 3-4 cell passages. Alternatively, a constitutive cell line can be maintained for up to three months by cell passaging; in this case protein expression takes five days.

### Protein purification

In both cases, isolated membranes were treated with a mild DDM/CHS detergent mixture to solubilize the target protein, followed with purification by metal-affinity chromatography. The major differences were in the cell lysis and membrane wash procedures.

In the case of Sf9, the cell pellet was lysed in a low osmotic buffer with tight douncing. Membranes were further purified with douncing in a high salt buffer to remove membrane-associated proteins. These procedures were repeated twice. Membranes were collected by centrifugation each time.

In the case of LEXSY, cells were disrupted in a microfluidizer with the addition of DNase I. The membrane fraction was as well collected by ultracentrifugation.

To increase stability in functional tests, the purified proteins from both expression systems were reconstituted into nanodiscs with MSP1D1 as a scaffold protein and POPC:POPG (7:3) lipid mixture. The protein-containing nanodiscs were further purified on a nickel-sepharose resin. The recovery of protein after reconstitution and purification in nanodiscs was 60% of the initial yield in micelles.

### Protein characterisation by SDS-PAGE, analytical SEC, and TSA

We assessed the purity, homogeneity, and thermostability of the obtained protein samples by SDS-PAGE, analytical SEC and TSA with the CPM (7-Diethylamino-3-(4’-Maleimidylphenyl)-4-Methylcoumarin) dye (Fig. 4 A-D). These methods are often applied to evaluate the quality of GPCR samples in structural studies ^28–32^.

**Figure 4.**
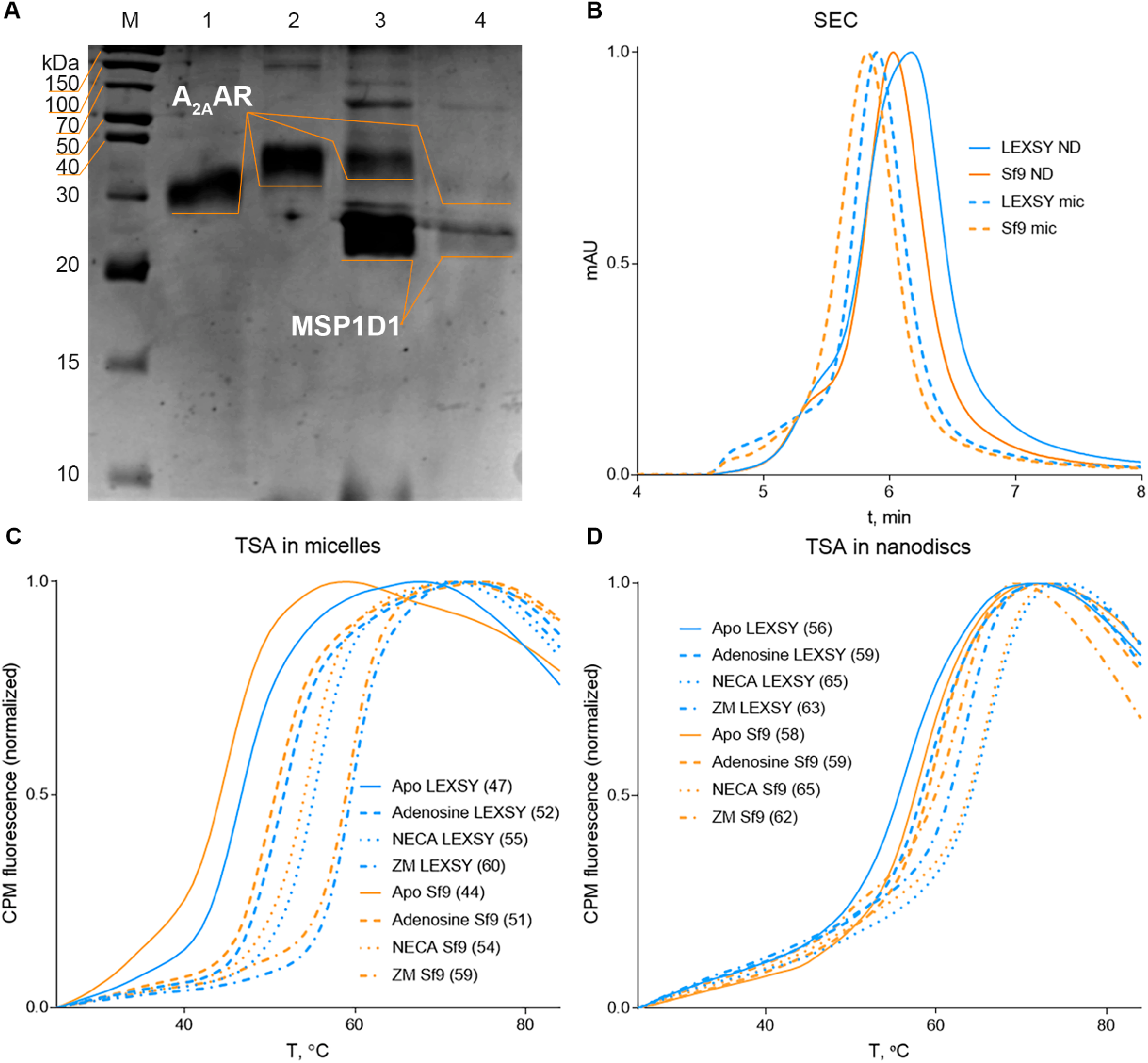
Characterization of the non-mutant A_2A_AR for the expression systems comparison. (**A**) SDS-PAGE analysis of the receptor expressed in Sf9 (lines 2,3) or LEXSY (lines 1,4) embedded in micelles (lines 1,2) and nanodiscs (lines 3,4). A_2A_AR (bands between 30-40 kDa) and MSP1D1 (bands between 20-30 kDa) are marked with orange lines. (**B**) Analytical SEC analysis of purified A_2A_AR from LEXSY and Sf9 as a complex with theophylline in micelles and apo in nanodiscs showing mostly monomeric protein preparations. (**C-D**) Thermal stability assay using CPM fluorescence of A_2A_AR stability in apo form, and in complexes with ligands: antagonist ZM 241385, and agonists NECA and adenosine, for the receptor in micelles (**C**) and nanodiscs (**D**). Melting temperature is shown in brackets.

SDS-PAGE showed the purity over 90% for A_2A_AR in micelles from both expression systems. For the receptors in nanodiscs, denaturing SDS-PAGE revealed bands for both MSP1D1 and A_2A_AR indicating on the successful nanodisc assembly (Fig. 4A).

Analytical SEC showed around 10% aggregation in the A_2A_AR samples in micelles and less than 20% aggregation in the A_2A_AR samples in nanodiscs, with a negligible difference between the expression systems. Monodispersity was assessed by the peak height and shape (Fig. 4B). The variation in mass, visualized by the SEC and SDS-PAGE, is explained by the difference in genetic constructs: 40 kDa for the Sf9 construct and 36.5 kDa for the LEXSY construct (Fig. 3).

TSA is based on the measurement of fluorescence from CPM dye that is enhanced when the dye interacts with the thiol and hydrophobic groups released upon heat-induced unfolding of the protein ^33,34^. The “melting temperature” is defined as the inflection point of the sigmoidal dependence of fluorescence on the sample temperature. TSA showed that in both membrane-mimicking environments and in two expression systems the stable protein sample was obtained (Fig. 4C-D): 47 °C and 44 °C in micelles, and 56 °C and 58 °C in nanodiscs for protein produced in LEXSY and Sf9, respectively. The stabilizing effects of ligands was measured as the ligand-induced increase of the melting temperature, indicating that protein was folded properly and was able to bind the ligands, the dispersion lied within the accuracy of the methodology, 2 °C ^33^: in micelles, 5 °C and 7 °C (adenosine), 8 °C and 10 °C (NECA), 13 °C and 15 °C (ZM 241385), and in nanodiscs, 3 °C and 1 °C (adenosine), 9 °C and 7 °C (NECA), 7 °C and 4 °C (ZM 241385) for LEXSY and Sf9 expression systems, respectively.

### smFRET for functional assessment

We used smFRET to test whether ligand binding to the extracellular side of the purified recombinant A_2A_AR translates into conformational changes on the intracellular surface of the receptor. In FRET, the energy transfer from blue-shifted donor to red-shifted acceptor fluorophores is sensitive to the inter-dye distance and orientation – thus, two fluorophores attached to the protein can report conformational changes that involve the attachment sites ^35^. To fluorescently label the receptor we introduced two cysteine mutations on the intracellular surface of the receptor: L225^6.27^C and Q310^8.65^C (Fig. 3), where the uppercase numbers indicate a helix with the position on it, according to the Ballesteros Weinstein numbering ^36^. We added two cysteine-reactive dyes (blue Alexa488-maleimide and red Atto643-maleimide) to the crude membranes containing the overexpressed receptor. We previously showed that this labeling scheme results in fluorescent labeling of the two newly introduced cysteins. The labeling was selective, as the native cysteines were shielded from dyes either by disulfide bonds or the lipid bilayer ^11^. We also found that smFRET using the double-labeled receptor was able to detect changes in protein conformation and dynamics induced by agonist. In the present work, the labeled receptors were purified and reconstituted into lipid nanodiscs as described above according to the scheme on Figure 2). Its quality was proved by SEC and TSA measurements (Supplementary Fig. 2).

We registered fluorescence of individual molecules in a diluted solution of the double-labeled A_2A_AR reconstituted in nanodiscs with a confocal fluorescence microscope as established previously ^11^ and described in Methods (“smFRET setup”). The obtained apparent FRET distributions (Fig. 5A) were very similar for A_2A_AR samples prepared using Sf9 and LEXSY expression systems. In both cases, agonist (NECA) binding increased FRET efficiency and shifted the peak of the apparent FRET distribution from ∼0.6 in apo and antagonist (ZM 241385)-bound receptor to ∼0.7 in agonist (NECA)-bound receptor (Supplementary Table 2).

**Figure 5.**
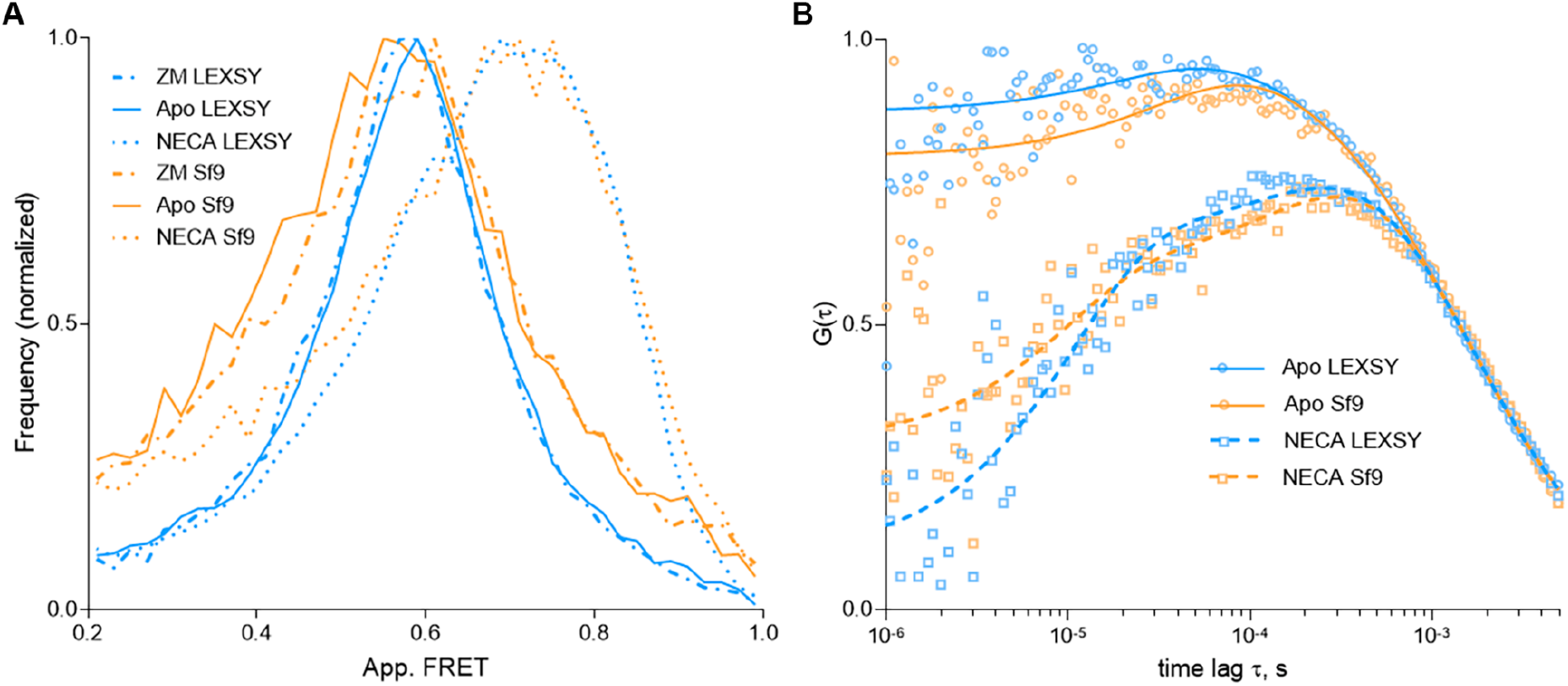
smFRET analysis for the expression systems comparison. (**A**) Apparent FRET distributions from smFRET experiments with nanodisc-reconstituted A_2A_AR expressed in Sf9 or LEXSY in apo form and in complexes with ligands: antagonist ZM 241385 and agonist NECA. (**B**) The cross-correlation fFCS function against time lag. Experimental points are shown as circles for the apo proteins and in squares for the complex with NECA. The fitting curves are plotted as solid lines for the apo proteins and as dotted for the complex with NECA.

Finally, we observed similar agonist-induced conformational dynamics in A_2A_AR expressed in LEXSY and Sf9 using filtered fluorescence correlation spectroscopy (fFCS) approach ^37^. fFCS curves for apo receptor deviate from the diffusion-related sigmoidal trend only on the 1–100 μs timescale, while for the agonist-bound receptors the deviation is also apparent on the 100–1000 μs timescale. As we showed previously ^11^, deviations on the shorter timescale are expected due to photoblinking of the fluorescent dyes, while deviations on the longer timescale are indicative of agonist-induced conformational dynamics of the receptor. We determined the exchange time of the agonist-induced dynamics *τ*_2_ = 0.26 ± 0.05 ms via fitting the experimental data. The data for A_2A_AR expressed in LEXSY and Sf9 were adequately described with χ _red_ ^2^ = 0.8 with *τ*_2_ globally linked between datasets (Supplementary Table 3). Our results revealed a consistent increase in protein dynamics upon agonist binding, which was demonstrated by an equivalent *A*_*2*_value across both LEXSY and Sf9 datasets.

## Discussion

A low recombinant expression of membrane proteins often represents a bottleneck for their structural and functional studies. To overcome it, one may need to apply various gene engineering strategies, such as modifications of the N- or C-termini or introduction of stabilizing mutations, as well as to try different orthologs of the target protein and expression enhancing additives ^38^. When choosing a suitable expression system one should take into account a number of factors: correct folding and functionality, post-translational modification pattern, potential proteolysis, expression yield, time, and cost. There is no single ideal approach that would work for all proteins, so it is important to expand the number of available expression systems.

Several expression systems have been established for GPCR structural studies including expression in mammalian, insect, yeast, and bacterial cells. For human GPCRs, expression in mammalian cell lines, such as human embryonic kidney (HEK) or Chinese hamster ovary (CHO), provides the closest-to-native conditions. For this reason, mammalian cell lines are widely used for functional cell-based assays of GPCR, however, their use in structural studies has been limited due to typically low protein yields and high production costs. By the end of 2022, 119 structures have been solved using GPCRs expressed in mammalian cell lines (Fig. 1, Supplementary Table 1).

Baculovirus expression system in insect cells is the most commonly used for structural research of GPCRs with 809 GPCR structures solved by the end of 2022. The advantages of expression in insect cells are eukaryotic expression and folding machinery, relatively high yields, and simplicity in comparison to mammalian systems. Its disadvantages include a high cost of growth medium, non-mammalian glycosylation patterns, distinct lipid membrane composition (e.g. lack of cholesterol), and cell lysis resulting from viral infection leading to the GPCR-containing cell membrane damage ^39^.

Yeast cell lines, e.g. *Pichia pastoris* and *Saccharomyces cerevisiae*, are suitable for overexpression of GPCRs and have the advantage of a fast growth rate. The downsides are reduced glycosylation, lipid composition different from mammalian cells ^40,41^ and challenges associated with rigidity of the yeast cell walls, which are difficult to disrupt during protein purification ^42^. Up to date, only two GPCRs expressed in *Pichia pastoris* were successfully used for structure determination: the histamine H1 receptor ^43^ and A_2A_AR ^17,18^.

Bacterial expression is the fastest, cheapest, and methodologically simplest, but the absence of eukaryotic machinery for post-translational modifications, including glycosylation, imposes serious limitations on functional folding of human GPCRs. Overexpressed GPCRs are frequently sorted into the inclusion bodies with the need for refolding, that is often a challenge ^44,45^. There have been only two human GPCR structures obtained using bacterial expression systems: a crystal structure of the alpha 1B adrenergic receptor ^46^ and an NMR-based structure of the chemokine receptor CXCR1 ^47^.

In this work, we reported a case study of expression, purification, and characterization of a model GPCR, A_2A_AR, in the LEXSY system. Leishmania can produce proteins in greater quantities compared to mammalian and insect cells. Typical expression levels in LEXSY range from 0.1 to 30 mg/L ^48^. In our case, the yield of A_2A_AR production in LEXSY was 3.6 mg/L, while the yield for Sf9 expression was 1 mg/L. The main advantages of the LEXSY approach include the ease of handling for cell cultures and fast growth rates (especially when compared to mammalian cells), the eukaryotic expression and folding machinery, the ability to up-scale expression by producing stable cell lines, the presence of post-translational modifications, including a glycosylation profile similar to mammalian proteins, resulting in functionally active folding of eukaryotic proteins, and a low cost. Cultivation is carried out on a simple inexpensive brain heart infusion medium with hemin supplementation.

For structural and functional studies of purified receptors, a high purity and monodispersity of the sample are required. We examined the purity of A_2A_AR samples via SDS-PAGE and assessed the aggregation state of the sample by SEC. We observed an over 90% purity for A_2A_AR samples in micelles produced in both expression systems as well as a successful nanodisc assembly. Over 80 % of monomeric fractions were demonstrated via SEC for A_2A_AR in both membrane-mimetic systems produced either in LEXSY or Sf9 expression systems.

An additional requirement for long room-temperature experiments, including crystallization trials and measurements of protein dynamics, is protein stability. Using TSA we showed that the thermal stability of A_2A_AR in the apo state as well as the stabilizing effects of ligands are the same regardless of the expression system used. This was demonstrated for A_2A_AR reconstituted in both micelles and nanodiscs.

Finally, we used smFRET to test whether the extracellular ligand binding translates into corresponding conformational changes and structural dynamics on the intracellular surface of the receptor. smFRET has been extensively used for investigation of protein structure and dynamics ^49^, and we recently showed that it can be used to study structural dynamics in A_2A_AR ^11,49^. The similar observed FRET efficiencies between samples produced in either LEXSY or Sf9 expression systems imply the similar final conformations of the intracellular part of the receptor. The shift of the apparent FRET distribution induced by the agonist binding confirms that conformational changes in the extracellular ligand binding pocket translates into structural changes on the intracellular surface of the receptor in both A_2A_AR samples. Moreover, fFCS revealed agonist-induced sub-millisecond conformational dynamics in A_2A_AR expressed in Sf9 and LEXSY. The exchange time for these conformational dynamics *τ*_2_ = 0.26 ±0. 05 ms is the same between the two expression systems and similar to the values reported previously ^11^.

In conclusion, we demonstrated that the A_2A_AR expressed in Sf9 and LEXSY shows similar monodispersity, stability in apo and ligand-bound states, agonist-induced conformational changes and structural dynamics. Thus, LEXSY has proven to be a well-suited expression system for recombinant production of functional receptors with a high yield, and therefore it could be added to the tool chest of common expression systems for GPCRs.

## Methods

### Materials

All used chemicals were of the purest grade available and were from Sigma–Aldrich with the exception of: X-tremeGENE™ HP DNA Transfection Reagent (Roche, Switzerland), Transfection Medium (Expression Systems, USA), Breathe-Easy membrane (Greiner BioOne, Austria), vent-cap flasks (Corning, USA), 4-(2-aminoethyl)benzenesulfonyl fluoride hydrochloride (AEBSF) (Gold Biotechnology, Argentina), Hemin (AppliChem, USA), penicillin (AppliChem, USA), streptomycin (AppliChem, USA), 6-aminohexanoic acid (6AHA) (AppliChem, USA), E-64 (Cayman Chemical, USA), leupeptin (Cayman Chemical, USA), aprotinin (AG Scientific, USA), protease inhibitor cocktail (cOmplete) (Roche, Switzerland), n-dodecyl-β-D-maltoside (DDM) (Anatrace, USA), Brain-Heart-Infusion Broth (Carl Roth, Germany), TALON resin (Clontech, USA), G-25 or PD-10 desalting column (GE Healthcare, USA), Ni-NTA resin (Qiagen, Germany), 7-Diethylamino-3-(4’-Maleimidylphenyl)-4-Methylcoumarin (CPM) (Invitrogen. USA), 1-palmitoyl-2-oleoyl-sn-glycero-3-phosphocholine (POPC) (Avanti Polar Lipids, USA), 1-palmitoyl-2-oleoyl-sn-glycero-3-phospho-(1’-rac-glycerol) (POPG) (Avanti Polar Lipids, USA), Atto643 maleimide (ATTO-TEC, Germany), Alexa488 maleimide (Invitrogen, USA), ZM 241385 (Cayman Chemical, USA), Nanofilm Sec 250 column (Sepax technologies, cat# 201250-4625, USA), Nunc Lab-Tek Chambered coverglass (Thermo Fisher Scientific, USA), NECA (Tocris, UK).

### Preparation of DNA construct for LEXSY expression

The nucleotide sequence encoding the human *ADORA2A* (1-316 aa) (UniProt ID C9JQD8) was codon optimized for *Leishmania tarentolae* expression by the GeneOptimizer software (Thermo Fisher Scientific, USA) and synthesized *de novo* (Eurofins, Luxembourg). The final gene construct was completed from the 3’-terminus by KpnI restriction site followed by 9×His-tag nucleotide sequence and inserted into the integrative inducible expression vector pLEXSY_I-blecherry3 (Jena Bioscience, Germany) via the BglII(5’) and NotI(3’) restriction sites. L225C^6.27^ and Q310C^8.65^ mutations were introduced by PCR.

Complete nucleotide sequence of A_2A_AR (1-316aa) in pLEXSY_I-blecherry3 plasmid, tag is shown in upper case:

atgccgattatgggtagcagcgtgtacatcacggtcgagctggcgattgcggtgctggcaattcttggcaacgtgctcgtgtgctgggccgtgtggcttaacagcaacctgcagaacgtgacgaactacttcgtggtgtctctggcggcagcggacattgcagtgggtgtcctggctatcccgttcgcgattacgatctcgacgggcttttgcgcagcatgccacggctgcctgtttatcgcatgcttcgtgctggtgctgacgcagagcagcatctttagcctgctcgcgatcgccatcgaccgctacattgcgattcgcatcccgctgcgctacaacggccttgtgactggtacacgcgcgaagggcatcattgcgatctgctgggtgctgagctttgcgattggcctgacaccgatgctcggctggaacaactgcggccagccaaaggagggcaagaaccattctcagggttgcggcgagggccaggtggcatgcctttttgaggatgtggtgccgatgaactacatggtgtacttcaactttttcgcgtgcgtgctggtcccgctgctgcttatgcttggcgtgtacctgcgcatcttcctggcagcacgtcgtcagctgaagcagatggagtctcagccacttccaggcgagcgtgcacgtagcacatgccagaaggaggtccacgcagctaagagcctggcgatcatcgtgggcctgtttgcgctttgctggctgccgctgcacatcatcaactgcttcacctttttctgcccggactgcagccacgcaccactgtggcttatgtacctggccatcgtgctgagccacacgaacagcgtggtcaacccgttcatctacgcgtaccgcatccgcgagttccgccagacgtttcgcaagatcattcgctcgcacgtgctgcgctgccaggagccttttaaggctggtaccCATCACCATCACCACCATCACCACCACtag

### Preparation of DNA construct for Sf9 expression

The nucleotide sequence encoding the human *ADORA2A* (2-316 aa) (UniProt ID C9JQD8) was obtained from the cDNA Resource Center (cdna.org, #ADRA2A0000) and modified with different tags for expression in *Spodoptera Frugiperda*. The final gene construct was completed from the 5’-terminus by nucleotide sequences of hemagglutinin signal peptide (MKTIIALSYIFCLVFA), FLAG-tag (DYKDDDDK), linker (AMGQPVGAP), and from the 3’-terminus by 10×His-tag nucleotide sequence and inserted into the pFastBac1 vector (Invitrogen, USA) via the BamHI(5’) and HindIII(3’) restriction sites. L225C^6.27^ and Q310C^8.65^ mutations were introduced by PCR.

Nucleotide sequence of A_2A_AR (2-316aa) in pFastBac1 plasmid, tags are shown in upper case: ATGAAGACGATCATCGCCCTGAGCTACATCTTCTGCCTGGTGTTCGCCGACTACAAGGACGATGATGACAAGGCCATGGGGCAACCCGTGGGCGCGCCAcccatcatgggctcctcggtgtacatcacggtggagctggccattgctgtgctggccatcctgggcaatgtgctggtgtgctgggccgtgtggctcaacagcaacctgcagaacgtcaccaactactttgtggtgtcactggcggcggccgacatcgcagtgggtgtgctcgccatcccctttgccatcaccatcagcaccgggttctgcgctgcctgccacggctgcctcttcattgcctgcttcgtcctggtcctcacgcagagctccatcttcagtctcctggccatcgccattgaccgctacattgccatccgcatcccgctccggtacaatggcttggtgaccggcacgagggctaagggcatcattgccatctgctgggtgctgtcgtttgccatcggcctgactcccatgctaggttggaacaactgcggtcagccaaaggagggcaagaaccactcccagggctgcggggagggccaagtggcctgtctctttgaggatgtggtccccatgaactacatggtgtacttcaacttctttgcctgtgtgctggtgcccctgctgctcatgctgggtgtctatttgcggatcttcctggcggcgcgacgacagctgaagcagatggagagccagcctctgccgggggagcgggcacggtccacactgcagaaggaggtccatgctgccaagtcactggccatcattgtggggctctttgccctctgctggctgcccctacacatcatcaactgcttcactttcttctgccccgactgcagccacgcccctctctggctcatgtacctggccatcgtcctctcccacaccaattcggttgtgaatcccttcatctacgcctaccgtatccgcgagttccgccagaccttccgcaagatcattcgcagccacgtcctgaggcagcaagaacctttcaaggcaCACCACCATCACCATCACCATCACCATCACtga

### LEXSY expression and membrane preparation

*Leishmania tarentolae* cells of the strain T7-TR (Jena Bioscience, Germany) were transfected with the A_2A_AR expression plasmids linearized by the SwaI restriction enzyme. After the clonal selection, the transfected cells were grown at 26 °C in the dark, with 120 rpm shaking at 2.5 cm orbital shaker in 2L, 5 mm-baffled flasks in the Brain-Heart-Infusion Broth supplemented with 5 μg/mL Hemin, 50 U/mL penicillin and 50 μg/mL streptomycin. When OD_600_=1 was reached, 10 μg/mL tetracycline was added, and incubation continued for an additional 24 h. The harvested cells (approximately 35 g of cells from 6L of culture) were resuspended in 150 ml of a buffer (50 mM HEPES, pH 7.0, 10 mM MgCl_2_, 20 mM KCl, 2 mM 6AHA), aliquoted and frozen by plunging in liquid nitrogen for storage at -80 °C until use.

For purification, cells were thawed on ice and disrupted in an M-110P Lab Homogenizer (Microfluidics, USA) at 10,000 psi in a lysis buffer (50 mM NaH_2_PO_4_/Na_2_HPO_4_, pH 7.6, 0.2 M NaCl, 20 mM KCl, 10 mM MgCl_2_, 10 % glycerol (w/v), 1 mM EDTA, 2 mM 6AHA, 50 mg/L DNase I and 1 tablet/50 mL cOmplete). The membrane fraction of the cell lysate was isolated by ultracentrifugation at 120,000×g for 1 h at 4 °C. The pellet was resuspended in the same buffer but without DNase I and stirred for 1 h at 4 °C. The ultracentrifugation step was repeated again.

### Sf9 expression and membrane preparation

High-titer recombinant baculovirus (>10^9^ viral particles per mL) for A_2A_AR expression in Sf9 insect cells was obtained following a modified Bac-to-Bac system protocol (Invitrogen, USA). Briefly, recombinant baculoviruses were generated by transfecting 1-5 μg of transfer bacmid into Sf9 cells (2.5 mL at a density of 10^6^ cells/mL) using 3 μL of X-tremeGENE™ HP DNA Transfection Reagent and 100 μL Transfection Medium. Cell suspension was incubated for 4 days with shaking using a Shel Lab incubator at 27 °C and 300 rpm in 24-deep well U-bottom plates covered with Breathe-Easy membrane. P0 viral stock was isolated by centrifugation at 2,000×g for 5 min, and used to produce high-titer baculovirus stock (P1): 40 ml of cells at 2 mln/mL density were infected with 2.5 mL supernatant, and grew for 72 h with shaking at 27 °C and 120 rpm (Innova 44, <u>New Brunswick</u>, Germany). Sf9 cells at a cell density of 2-3 ×10^6^ cells/mL were infected with P1 virus at multiplicity of infection equal to 5. Expression was performed with shaking at 27 °C and 120 rpm (Innova 44, <u>New Brunswick</u>, Germany), in 1 L vent-cap flasks. Cells were harvested by centrifugation at 2,000×g for 10 min, after 48 h post infection, and stored at -80 °C until further use. Cell count, viral titers and expression level were performed by flow cytometry on BD Accuri C6 (BD Biosciences, USA).

The biomass obtained from 250 mL of cell culture was thawed at 4 °C in 70 mL of low-salt buffer (10 mM HEPES pH 7.5, 10 mM MgCl_2_, 20 mM KCl) (scaled if necessary) supplemented with 50 μL of a protease inhibitor cocktail (500 μM AEBSF, 1 μM E-64, 1 μM leupeptin, 150 nM aprotinin). The mixture was homogenized in a high-tight 100 mL Potter douncer at 4 °C, centrifuged for 20 min at 4 °C and 220,000×g. The supernatant was discarded, and the pellet was resuspended with douncer in 70 mL of high-salt buffer (10 mM HEPES, pH 7.5, 10 mM MgCl_2_, 20 mM KCl, 1 M NaCl) supplemented with 50 μL of the protease inhibitor cocktail and centrifuged for 30 min at 4 °C and 220,000×g. The last step was repeated twice.

### Solubilization and purification of recombinant A_2A_AR

All procedures were performed at 4 °C. The washed membranes from 250 mL of cell culture biomass (both from LEXSY and Sf9) were homogenized in 12.5 mL of a resuspension buffer (50 mM HEPES, pH 7.5, 10 mM MgCl_2_, 20 mM KCl, 800 mM NaCl, 4 mM theophylline, 2 mg/mL iodoacetamide, 10% w/v glycerol, 50 μL protease inhibitor cocktail). The suspension was incubated for 30 min in the dark. Then 12.5 mL of a solubilization buffer (2/0.4% w/v DDM/CHS, 50 mM HEPES, pH 7.5, 10 % w/v glycerol, 800 mM NaCl) was added. Solubilization was carried out for 3 h with constant rotation on an orbital shaker (15 rpm); the insoluble debris was eliminated then by centrifugation for 1 h at 650,000×g, while the target protein remained in the supernatant. The supernatant was added to 250 μl of TALON resin equilibrated with an application buffer (50 mM HEPES, pH 7.5, 10 % w/v glycerol, 800 mM NaCl) and the mixture was incubated overnight with constant rotation (15 rpm).

The next day, the resin was deposited into a gravity column, washed with 10 column volume (CV) of wash buffer 1 (0.1/0.02% DDM/CHS,100 mM HEPES pH 7.5, 10% w/v glycerol, 800 mM NaCl, 4 mM theophylline, 10 mM MgCl_2_, 25 mM imidazole, 8 mM ATP, 25 μL protease inhibitor cocktail), then 10 CV of wash buffer 2 (0.05/0.01% w/v DDM/CHS, 25 mM HEPES, pH 7.5, 10% w/v volume glycerol, 800 mM NaCl, 4 mM theophylline, 50 mM imidazole, 25 μL protease inhibitor cocktail). The protein was eluted in the elution buffer (0.025/0.005% w/v DDM/CHS, 25 mM HEPES, pH 7.5, 10% w/v glycerol, 800 mM NaCl, 4 mM theophylline, 10 μL protease inhibitor cocktail, 220 mM imidazole), and then desalted from imidazole using size-exclusion PD-10 column equilibrated with desalt buffer (0.025/0.005% w/v DDM/CHS, 25 mM HEPES, pH 7.5, 800 mM NaCl). The protein yield was estimated by light absorption at 280 nm using the Biotek Synergy H4 plate reader equipped with the Take3 microvolume plate.

### Nanodisc assembly

The Membrane Scaffold Protein 1D1 (MSP1D1) was expressed in *E*.*coli* using a gene with an N-terminal 6×His-tag and an upstream TEV-protease site cloned into pET28a(+) (Addgene plasmid #20061). MSP1D1 was purified using IMAC with further cleavage of 6×His-tag by TEV protease ^50,51^. The lipid mixture of POPC:POPG in chloroform was prepared at a molar ratio 7:3. The lipid film was dried under a gentle nitrogen stream, followed by removal of the solvent traces under vacuum, and then was solubilized in 100 mM sodium cholate. The purified A_2A_AR in DDM/CHS micelles was mixed with MSP1D1 and the POPC:POPG lipid mixture at a molar ratio A_2A_AR:MSP1D1:lipids=0.2:1:60. The final sodium cholate concentration was adjusted to 25 mM, the typical final receptor concentration was no less than 0.1 mg/mL. After 1 h incubation at 4 °C, the mixture was incubated with wet Bio-Beads SM-2 (0.14 g of beads for 1 g of detergent were washed in methanol and equilibrated with 25 mM HEPES, pH 7.5, 800 mM NaCl) overnight at 4 °C with constant rotation on an orbital shaker. The next morning, a fresh portion of Bio-Beads for an additional 4 h incubation was added, beads were discarded then and the supernatant containing reconstituted into nanodiscs A_2A_AR was incubated with 250 μL of Ni-NTA resin (Qiagen, Germany) (2 h with constant rotation) for separating from empty nanodiscs. The protein was eluted in the elution buffer (25 mM HEPES, pH 7.5, 800 mM NaCl, 220 mM imidazole), and then desalted from imidazole using a size-exclusion PD-10 column equilibrated with desalt buffer (25 mM HEPES, pH 7.5, 150 mM NaCl). The protein yield is estimated by light absorption at 280 nm using the Biotek Synergy H4 plate reader equipped with the Take3 microvolume plate (Agilent, USA).

### SDS-PAGE and size-exclusion chromatography (SEC)

The samples were subjected to SDS-PAGE using a Mini Protean Tetra system (Bio-Rad, USA) and agarose gel (5% concentrating gel with AA:bisAA ratio of 29:1 and 15% resolving gel with 19:1 AA:bisAA ratio). 5 μg of receptor per lane premixed with a loading buffer (25 mM Tris, pH 6.8, 25 % glycerol, 0.25 % SDS, bromophenol blue) was loaded in the without preheating, and the gel was stained after separation with Coomassie Brilliant Blue R-250.

Analytical size-exclusion chromatography was performed on a Dionex Ultimate 3000 instrument (<u>Thermo Fisher Scientific</u>, USA) equipped with a Nanofilm Sec 250 gel filtration analytical column. The column was equilibrated with a chromatographic buffer: 0.05/0.01% w/v DDM/CHS, 25 mM HEPES, pH 7.5, 500 mM NaCl, 20 mM MgCl_2_, 2% w/v glycerol. For nanodiscs no detergents were added. The flow rate was 0.35 mL/min, protein absorption was detected at 280 nm, and 40 μL of the sample was injected.

### Thermal stability assay (TSA)

TSA measurements were carried out on a Rotor-Gene Q 6 plex (Qiagen, Germany) instrument at a heating rate of 2 °C/min and a temperature range of 25−90 °C in the presence of a CPM dye. The excitation wavelength was set at 387 nm and the emission wavelength was 463 nm. The A_2A_AR concentration was about 2 μM. Buffer conditions: 0.025/0.005% w/v DDM/CHS, 25 mM HEPES, pH 7.5, 800 mM NaCl for the micelles, and 25 mM HEPES, pH 7.5, 150 mM NaCl for nanodiscs. To obtain a good fluorescent intensity we used a 2.5-fold molar excess of the CPM dye to protein. To prepare protein for the ligand-binding measurements we added 200 μM of ZM 241385 or NECA and incubated for 1 h at 4 °C in the dark. The TSA was performed in a total volume of 50 μL. Protein melting temperatures were defined using the GraphPad Prism v.7 fitted to the Boltzmann sigmoid function.

### Labeling

The washed membranes containing the double-mutant A_2A_AR (L225^6.27^C/Q310^8.65^C) were resuspended in a labeling buffer (50 mM HEPES, pH 7.0 10 mM MgCl_2_, 20 mM KCl, 2 mM 6AHA, and 1 tablet/50 mL cOmplete) and mixed with Atto643 maleimide and Alexa488 maleimide, dissolved in DMSO (0.5 mg of each fluorescent dye per 10 g of cells). The labeling reaction was carried out overnight in the dark at 4 °C with a constant rotation on an orbital shaker.

The next day, the membrane fractions were pelleted by ultracentrifugation at 120,000×g for 1 h at 4 °C and washed twice with the labeling buffer to remove the unbound fluorescent labels.

### smFRET setup

For single-molecule experiments, a home-built multi-parameter fluorescence detection microscope with pulsed interleaved excitation (MFD PIE) was used as described previously ^11^. The inverted microscope stand (IX70, Olympus Belgium NV, Berchem, Belgium) was equipped with a 60x water-immersion objective (NA=1.2, UPLSAPO-60XW, Olympus). Two spatially overlapped lasers were used to excite donor and acceptor fluorophores: a pulsed 483-nm laser diode (LDH-P-C-470, Picoquant) and a pulsed 635-nm laser diode (LDH-P-C-635B, Picoquant, Germany), with alternating at 26.67 MHz pulses, delayed by 18 ns with respect to each other. Sample emission was transmitted through a pinhole and spectrally split. Both the blue range and red range were split by polarization into two detection channels. Photons were detected by four avalanche photodiodes (PerkinElmer SPCM-AQR12/14 and Laser Components COUNT BLUE): B_||_ (blue-parallel), B_⊥_ (blue-perpendicular), R_||_ (red-parallel) and R_⊥_ (red-perpendicular), which were connected to a TCSPC device (SPC-630, Becker & Hickl GmbH, Germany). Microscope alignment (excitation light guiding, objective lens correction collar, pinhole, detectors) was done using real-time fluorescence correlation spectroscopy (FCS) on freely diffusing Atto_488_-COOH and Atto_655_-COOH in water. For more details about the used equipment the reader is referred to ^11^.

### smFRET experiment

Samples of double-labeled A_2A_AR in nanodiscs were diluted in a buffer, containing 20 mM HEPES, pH 7.5, 150 mM NaCl to a protein concentration of 0.5-2 nM. To measure the effects of ligand binding, samples were supplemented with either 10 μM ZM 241385 or 10 μM NECA and incubated for 30 min at 4 °C. After incubation, the samples were transferred to a Nunc Lab-Tek Chambered coverglass (Thermo Fisher Scientific, USA). smFRET experiments were performed at 100 μW of 483 nm and 50 μW of 635 nm excitation. Measurements were recorded at room temperature (22 °C), samples were replenished every 30 min. With all filters applied (see “smFRET data processing” in Methods), 11,000-24,000 fluorescence bursts corresponding to double-labeled molecules were collected for each sample: 11,946 bursts for apo A_2A_AR from Sf9, 11,501 and 16,311 bursts for ZM241385- and NECA-bound A_2A_AR from Sf9, respectively; 21,721 bursts for apo A_2A_AR from LEXSY, 19,803 and 23,920 bursts for ZM 241385- and NECA-bound A_2A_AR from LEXSY. Background scattering information was obtained via a buffer measurement under identical conditions.

### smFRET data processing

For single-molecule data, we employed a two-color MFD all-photon burst search algorithm ^52^ using a 500 μs sliding time window (min. 50 photons per burst, min. 5 photons per time window). A 0-20-ms burst duration cutoff was applied to remove sparse (< 1%) slow moving aggregates. The absolute burst-averaged apparent FRET efficiency and stoichiometry S_raw_ were calculated as:

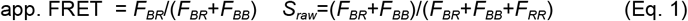

where *F*_*BB*_ and *F*_*BR*_ is the background-corrected number of photons in the blue and red detection channels after blue excitation, respectively, and *F*_*RR*_ is the background-corrected number of photons in the red detection channels after red excitation (independently of the polarization in all three cases) ^53^.

Using the stoichiometry *S*_*raw*_, we selected molecules labeled with both donor and acceptor fluorophores for further smFRET analysis and distinguish them from single-labeled molecules that also originate from our stochastic labeling procedure. Molecules labeled with donor dye fluoresce after the pulses of blue laser: the direct emission of the donor contributes to the donor color channel (*F*_*BB*_) and FRET signal, if present, contributes to the acceptor color channel (*F*_*BR*_). Molecules labeled with acceptor dye fluoresce after pulses of the red laser (*F*_*RR*_). To select double-labeled subpopulations of molecules, we used specific restrictions for the stoichiometry S_raw_, and kernel-density estimator ALEX-2CDE ^54^: ALEX-2CDE < 15, 0.2 < *S*_raw_ < 0.5 (Supplementary figure 3).

All analyses of experimental smFRET data were performed in the software package PAM (PIE Analysis with MATLAB) ^55^.

### Filtered fluorescence correlation spectroscopy (fFCS)

fFCS analysis was described in detail in our recent study on structural dynamics of A_2A_AR ^11^. The mathematical background of fFCS was previously described in detail ^56^. Here we used the same procedure with minor adjustments described below.

In short, we built two reference TCSPC patterns corresponding to the “low-FRET” pseudo-species 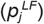 and “high-FRET” pseudo-species 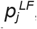. For this, we merged four smFRET datasets for A_2A_ AR with apo or NECA-bound receptors expressed in LEXSY or Sf9; bursts corresponding to double-labeled receptors with 0.1 < *app. FRET* < *0*.*3* were used to build *p*_j_ ^*LF*^, bursts corresponding to double-labeled receptors with 0.8<*app. FRET* <*0*.*9* were used to build 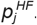. Four TCSPC channels for photons in parallel and perpendicular polarizations and in two color channels (emission of donor and FRET-sensitized emission of acceptor, both after donor excitation) were stacked into a single array and indexed with *j* for global analysis. Using these reference TCSPC patterns filters 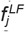 and 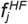 were calculated as described in ^56^. To reduce noise in fFCS filters, at this step TCSPC bin was increased to 100 μs.

Using the reference filters 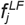 and 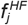 and the fluorescence signal *S*_*j*_, the correlation functions *G(τ)* were calculated for each dataset:

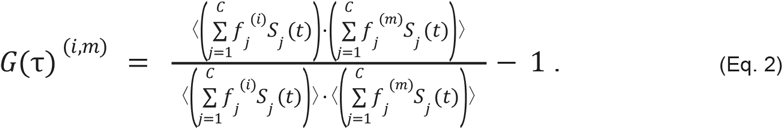

Only bursts from double-labeled molecules were taken into account; a 10-ms time window was introduced to reduce artifacts related to the sub-ensemble FCS analysis. The cross-correlation functions G^(HF,LF)^ were fit using equation:

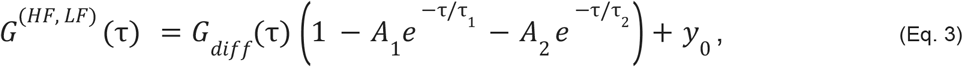

where the diffusion-limited term is:

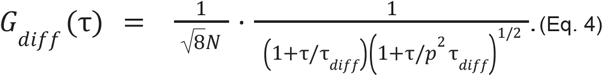

We fit fFCS curves with one anticorrelating term for the apo A_2A_AR (*A*_*1*_>0, *A*_*2*_=0) and two anticorrelating terms (*A*_*1*_>0, *A*_*2*_>0) for the agonist-bound receptor. The fast anticorrelation term (*A*_*1*_) was assigned to dye photophysics; the slow anticorrelation term (*A*_2_) represents agonist-induced conformational dynamics of A_2A_ AR. Diffusion time τ_*diff*_ and relaxation time of the agonist-induced conformational dynamics τ_2_ were globally optimized across all datasets.

The resulting cross-correlation curves were normalized using *N* and offset *y*_*0*_ and plotted in Fig. 5B. The 95% confidence intervals for the fitting parameters were calculated using the numerical Jacobian matrix.

## Acknowledgements

Authors thank Taras Balandin for the advising and discussion during project realization.

This work is supported by RSCF research grant 22-74-00024. J.H. acknowledges support from the Flemish government through long-term structural funding Methusalem (CASAS2, Meth/15/04).

## Author contributions statement

P.K., A.L., M.S. expressed protein in Sf9 and prepared membranes under supervision of A.M. and V.C.

O.V. expressed protein in LEXSY under supervision of V.I.

D.K., E.B. prepared LEXSY membranes under supervision of P.Ku.

P.K., I.M., O.V., M.S., A.B., D.D., A.L. solubilized, purified and reconstructed protein in nanodiscs.

P.K., A.B., D.D. performed SDS-PAGE, SEC and TSA under guidance of A.M. and V.C.

I.M., O.V., A.B. and P.K. performed labeling.

I.M. collected smFRET data and analyzed data together with J.Ho., J.H., T.G., and V.B.

A.K. analyzed data on expression systems of available structures.

A.L. produced the first draft of the manuscript. V.B., I.M., P.Kh. and A.L. conceptualized the study.

V.B. supervised the work.

All the authors contributed to analyzing data, writing the original draft, reviewing, and editing.

## Ethics declarations

Competing interests. The authors declare no competing interests

