## Supplementary information for "Functional GPCR expression in eukaryotic LEXSY system"

### Supplementary information
### for
### Functional GPCR expression in eukaryotic LEXSY system

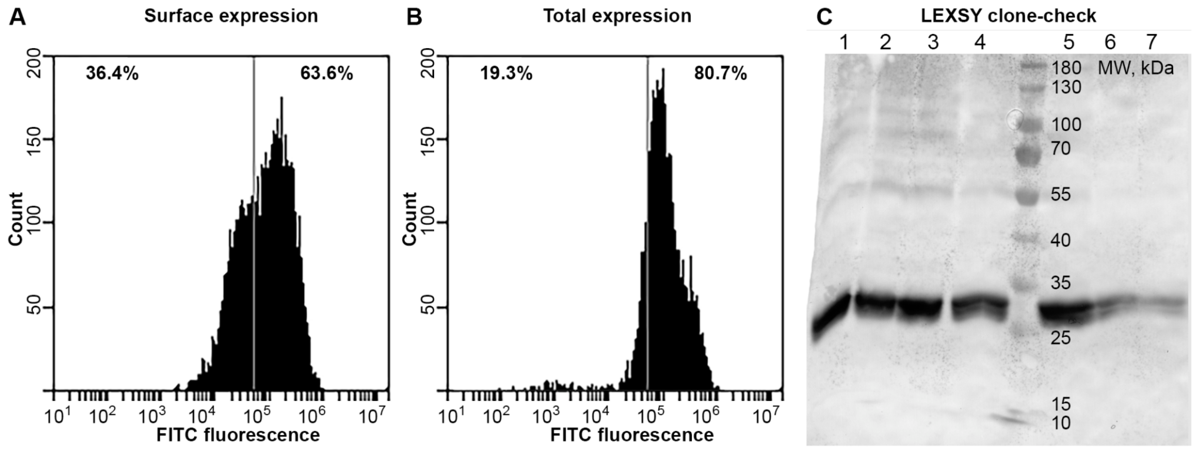

**Figure S1. Expression analysis for both systems.**
Flow cytometry analysis of the surface (**A**) and total (**B**) expression in Sf9 cells (day 14 of the Sf9 pipeline). Expression was measured on the BD Accuri C6 cytometer with the addition of anti-FLAG FITC-conjugated antibodies. Total expression was measured by addition of Triton X-100. FITC fluorescence intensity was measured in relative units. Non-infected cells treated with the same antibodies were used for the threshold establishing. Western blotting analysis of the LEXSY clones (**C**). Clone #3 was chosen for the further biomass production.

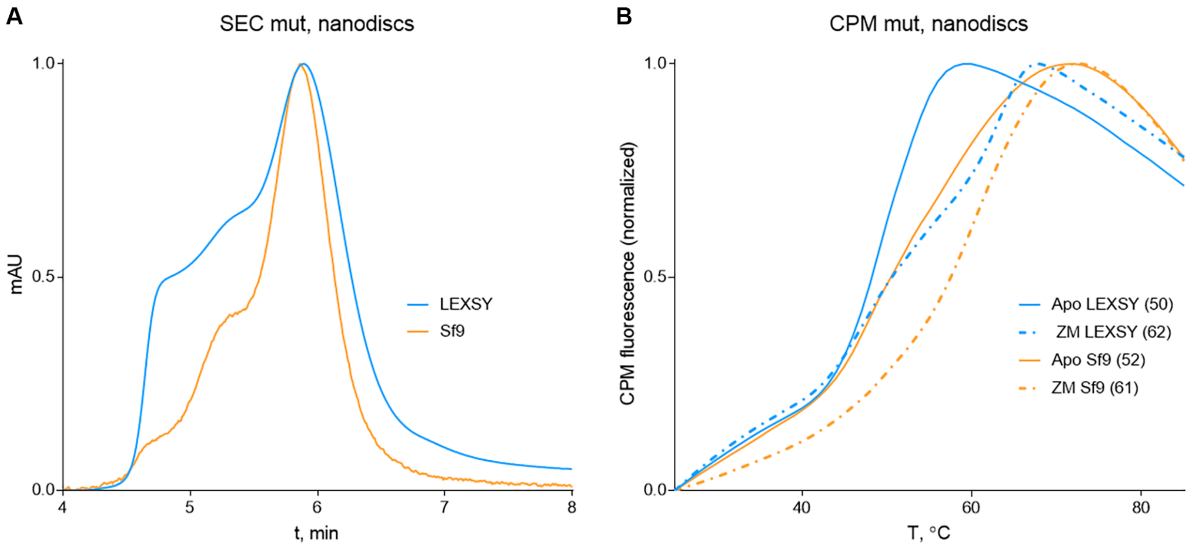

**Figure S2. Characterization of the labeled double-mutant A2AAR (L225$^{6.27}$C/Q310$^{8.65}$C).**
(**A**) Analytical SEC analysis of purified $A_{2A}$AR in nanodiscs. (**B**) Thermal stability assay of $A_{2A}$AR stability in apo form, and in complex with ZM241385 for the receptor in nanodiscs. Melting temperature is shown in brackets.

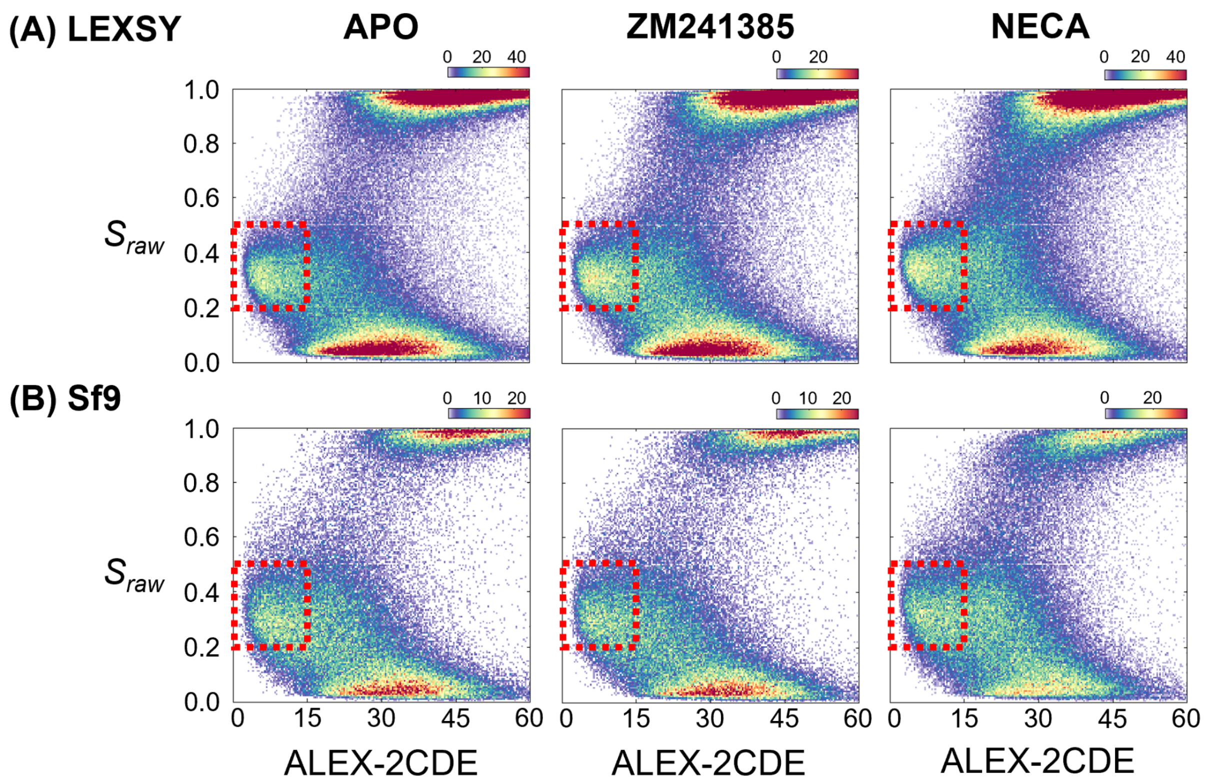

**Figure S3. Selection of double-labeled $A_{2A}$AR molecules using ALEX-2CDE and $S_{raw}$ filters.**
Two-dimensional distribution of $S_{raw}$ and ALEX-2CDE values for fluorescence bursts registered in smFRET experiment with $A_{2A}$AR (**A**) expressed in LEXSY or (**B**) expressed in Sf9. Red rectangles show the regions corresponding to double-labeled molecules selected with filters $0.2<S_{raw}<0.5$ and ALEX-2CDE<15.

**Table S1. The number of GPCR structures obtained using various heterologous expression systems between 2007 and 2022.**
The samples produced through homologous expression are not included. In total, 948 structures were analyzed, with corresponding PDB IDs provided The data are based on the updated version of the GPCRdb [1] database (as of March 9, 2023) and cross-referenced with RCSB PDB [2,3] for information on the expression system. In some rare cases, PDB ID-related articles were manually analyzed to determine the expression system.

| System / Year | Mammalian | Insect | Bacterial | Yeast |
|---|---|---|---|---|
| **2007** | **1** | **3** | **0** | **0** |
| | 2J4Y | 2RH1, 2R4S, 2R4R | | |
| **2008** | **0** | **3** | **0** | **0** |
| | | 3D4S, 2VT4, 3EML | | |
| **2009** | **0** | **0** | **0** | **0** |
| **2010** | **0** | **10** | **0** | **0** |
| | | 3KJ6, 3NYA, 3NY9, 3NY8, 3OE8, 3OE9, 3OE6, 3OE0, 3ODU, 3PBL | | |
| **2011** | **1** | **18** | **0** | **1** |
| | 2X72 | 3PDS, 2Y04, 2Y03, 2Y02, 2Y00, 3P0G, 3QAK, 2Y01, 2YDO, 2YDV, 2YCZ, 2YCX, 2YCW, 2YCY, 3SN6, 3PWH, 3REY, 3RFM | | 3RZE |
| **2012** | **1** | **16** | **0** | **2** |
| | 4A4M | 3UON, 3V2W, 3V2Y, 4DAJ, 4DKL, 4DJH, 3UZC, 3UZA, 4EA3, 4EJ4, 4AMI, 4AMJ, 4EIY, 4GRV, 4GBR, 3VW7 | | 3VG9, 3VGA |
| **2013** | **2** | **17** | **0** | **0** |
| | 4BEZ, 4BEY | 4GPO, 4IB4, 4IAR, 4IAQ, 3ZPQ, 3ZPR, 4JKV, 4K5Y, 4L6R, 4MBS, 4LDL, 4LDO, 4LDE, 4MQT, 4MQS, 4NC3, 4N6H | | |
| **2014** | **0** | **15** | **4** | **0** |
| | | 4N4W, 4O9R, 4OR2, 4NTJ, 4BVN, 4PXZ, 4PY0, 4OO9, 4PHU, 4QIM, 4QIN, 4QKX, 4U15, 4U14, 4U16 | 4BUO, 3ZEV, 4BV0, 4BWB | |
| **2015** | **3** | **22** | **0** | **0** |
| | 4XT3, 4XT1, 4ZWJ | 4S0V, 4RWD, 4RWA, 4RWS, 4XNW, 4XNV, 4UHR, 4UG2, 4YAY, 4Z35, 4Z36, 4Z34, 4XES, 4XEE, 5C1M, 5CGD, 5CGC, 5A8E, 4ZUD, 5DHH, 5DHG, 5F8U | | |
| **2016** | **6** | **26** | **0** | **0** |
| | 5DGY, 5L7D, 5L7I, 5EN0, 5DYS, 5TGZ | 5D5A, 5D5B, 4ZJ8, 4ZJC, 5CXV, 5DSG, 5EE7, 4Z9G, 5IUA, 5IU8, 5IUB, 5IU7, 5IU4, 5JQH, 5G53, 5D6L, 5GLH, 5GLI, 5K2D, 5K2A, 5K2B, 5K2C, 5U09, 5LWE, 5T1A, 5T04 | | |
| **2017** | **5** | **39** | **0** | **0** |
| | 5V57, 5V56, 5XRA, 5XR8, 5W0P | 5TVN, 5UIG, 5UEN, 5UNH, 5UNG, 5UNF, 5NDD, 5NJ6, 5NDZ, 5UZ7, 5VEX, 5XF1, 5XEZ, 5VEW, 5VAI, 5UVI, 5VBL, 5JTB, 5TZY, 5TZR, 5NX2, 5UIW, 5N2R, 5MZJ, 5MZP, 5N2S, 5TUD, 5XSZ, 5X93, 5XPR, 5X7D, 5NLX, 5NM2, 5NM4, 5WIU, 5WIV, 5WQC, 5WS3, 5VRA | | |
| **2018** | **13** | **67** | **0** | **2** |
| | 6FKD, 6FK9, 6FK6, 6FKC, 6FKB, 6FK8, 6FKA, 6FK7, 5WB2, 5WB1, 6FUF, 6MEO, 6MET | 5O9H, 6B73, 5YQZ, 5OLV, 5OLZ, 5OLG, 5OM4, 5OLO, 5OM1, 5OLH, 5V54, 6BQH, 6BQG, 6B3J, 5WF5, 5WF6, 6FFH, 6FFI, 6CM4, 5ZBQ, 5ZBH, 5KW2, 6GDG, 6D35, 6D32, 6C1Q, 6C1R, 6DDF, 6DDE, 5ZKQ, 5ZKP, 6CMO, 6D9H, 6G79, 5XJM, 6BD4, 6DRY, 6DRZ, 6DRX, 6DS0, 6E3Y, 6D26, 6D27, 6H7O, 6H7L, 6H7N, 6H7J, 6H7M, 6AKY, 6AKX, 6MXT, 6FJ3, 6IGL, 6IGK, 5ZKC, 5ZKB, 5YC8, 5ZK8, 5ZK3, 5ZHP, 5YHL, 5YWY, 6AK3, 6M9T, 6E59, 6IIV, 6IIU | | 5X33, 6AQF |
| **2019** | **6** | **67** | **0** | **0** |
| | 6DO1, 6OT0, 6O3C, 6I9K, 6QNO, 6KUY | 6GPX, 6GPS, 6IBL, 6HLP, 6HLL, 6HLO, 6N52, 6N51, 6NIY, 5ZTY, 6N4B, 6A94, 6A93, 6J21, 6J20, 6NBH, 6NBI, 6NBF, 6ME8, 6ME6, 6ME9, 6ME7, 6ME4, 6ME3, 6ME2, 6ME5, 6MH8, 6OIK, 6OIJ, 6E67, 6N48, 6GT3, 6OS9, 6OSA, 6K1Q, 6IBB, 6RNK, 6QZH, 6KQI, 6RZ4, 6RZ5, 6JZH, 6PS8, 6KK7, 6KJV, 6KK1, 6PS6, 6PS4, 6PS2, 6PS0, 6PS5, 6PS3, 6PS1, 6PRZ, 6PS7, 6NI3, 6PWC, 6IQL, 6KUW, 6KUX, 6PT2, 6PT3, 6RZ9, 6RZ7, 6RZ8, 6RZ6, 6OL9 | | |
| **2020** | **18** | **105** | **0** | **0** |

| | | | | |
|---|---|---|---|---|
| | 6KPG, 6OS2, 6OS1, 6OS0, 6U1N, 6LPB, 6LW5, 6WIV, 7C7S, 7C7Q, 6WW2, 6XBL, 6XBK, 6XBM, 6XBJ, 7CUM, 7CA5, 7CA3 | 6TPJ, 6TPG, 6TPN, 6TQ7, 6TQ9, 6TQ6, 6TP4, 6TOD, 6TQ4, 6TP6, 6TP3, 6TO7, 6ORV, 6TOS, 6TOT, 6JOD, 6KNM, 6P9Y, 6P9X, 6LRY, 6PB1, 6PB0, 6PT0, 6KPF, 6KPC, 6UP7, 6LI2, 6LI1, 6LI3, 6LI0, 6OMM, 6LUQ, 6M1H, 6M1I, 6KP6, 6VI4, 6LN2, 6UUN, 6OBA, 6LML, 6LMK, 6UVA, 6UUS, 6K42, 6K41, 6W25, 6WHC, 6VJM, 6UOA, 6UO9, 6UO8, 6VMS, 6TKO, 6WWZ, 6W2X, 6W2Y, 6WJC, 6V9S, 6S0L, 6S0Q, 6VCB, 7C6A, 7C61, 6TPK, 7BZ2, 6WZG, 6WI9, 6WPW, 6KO5, 7C2E, 6VN7, 7BW0, 6LFO, 6LFM, 6LFL, 7JJO, 7CFM, 7CFN, 6X19, 6X1A, 6X18, 6Z10, 6ZDR, 6ZDV, 6WH4, 6WHA, 6WGT, 7BR3, 7D3S, 7D7M, 6XOX, 7CZ5, 6WQA, 6LPL, 6LPJ, 6LPK, 7BVQ, 7BU6, 7BU7, 7BTS, 7AD3, 7D68, 7DHR, 7DHI, 7DFP | | |
| **2021** | **27** | **143** | **12** | **0** |
| | 7L1U, 7L1V, 7EB2, 7KH0, 7DD5, 7DD7, 7DD6, 7EPC, 7EPA, 7EPD, 7EPB, 7JHJ, 7DB6, 7F9Y, 7FD9, 7FD8, 7E6U, 7E6T, 7RMI, 7RMG, 7RMH, 7V3Z, 7EWR, 7EWP, 7SHF, 7EWL, 7SHE | 7LCI, 7LCK, 7LCJ, 7DDZ, 7D77, 7D76, 7K15, 7JVR, 7JVP, 7JVQ, 7JV5, 7KNU, 7KNT, 7LJD, 7CRH, 7CKY, 7CKW, 7LJC, 7CKZ, 7CKX, 7CMV, 7CMU, 7DTW, 7DTU, 7DTV, 7DTT, 7DFL, 7ARO, 7JOZ, 7E33, 7E2Z, 7E2X, 7E2Y, 7E32, 7AUE, 7CX3, 7CX4, 7CX2, 7MBY, 7MBX, 7BB7, 7BB6, 7C4S, 7E9H, 7EPE, 7E9G, 7EPF, 7O7F, 7M3G, 7M3E, 7M3J, 7M3F, 7MTR, 7MTS, 7MTQ, 7E14, 7DUQ, 7F1T, 7F1S, 7F1R, 7F1Q, 7DW9, 7KI1, 7KI0, 7DTY, 7EVW, 7DH5, 7EXD, 7EVM, 7DUR, 7F16, 7F9Z, 7M8W, 7EZM, 7EZK, 7EZH, 7F4I, 7F4H, 7F4F, 7F4D, 7P2L, 7RM5, 7LD4, 7LD3, 7DGE, 7DGD, 7EW1, 7EW4, 7EW2, 7EW3, 7EW7, 7EW0, 7EVZ, 7EVY, 7FII, 7FIJ, 7FIH, 7FIG, 7RTB, 6ZFZ, 6ZG4, 6ZG9, 7F8V, 7F8W, 7F8Y, 7F8U, 7F2O, 7EIB, 7V9M, 7F53, 7F54, 7F58, 7F55, 7S8P, 7S8M, 7S8N, 7S8L, 7S8O, 7PIU, 7PIV, 7S0F, 7S0G, 7VV6, 7VV5, 7VV4, 7VV3, 7VV0, 7VUZ, 7VUY, 7VDM, 7VDL, 7VDH, 7P02, 7P00, 7NA8, 7NA7, 7VOE, 7VOD, 7VUJ, 7VUH, 7VUI, 7VUG, 7F8X | 7L0R, 7L0P, 7L0S, 7L0Q, 6ZA8, 6Z4V, 6Z4S, 6Z4Q, 6ZIN, 6Z8N, 6Z66, 6YVR | |
| **2022** | **36** | **256** | **1** | **0** |
| | 7SIM, 7SIN, 7SIL, 7RKN, 7RKM, 7RKF, 7RYC, 7T6B, 7TUZ, 7TUY, 7ZI0, 7F23, 7F1Z, 7F24, 7F0T, 7X2D, 7F1O, 7X2F, 7XJL, 7XJK, 7X2C, 7UTZ, 7T9N, 7T9I, 8CXO, 7VVO, 7VVN, 7VVM, 7VVL, 7VVK, 7VVJ, 7T9M, 7T32, 7Y27, 7Y26, 7Y24 | 7WF7, 7EO4, 7EO2, 7S3I, 7S1M, 7F6H, 7F6I, 7LLY, 7LLL, 7W2Z, 7F83, 7WC9, 7WC8, 7WC7, 7WC5, 7WC6, 7WC4, 7TD4, 7TD3, 7TD2, 7TD1, 7TD0, 7JNI, 7VGX, 7FIN, 7VH0, 7VGY, 7VGZ, 7V35, 7VBI, 7FIM, 7VAB, 7FIY, 7PYR, 7PX4, 7T11, 7T10, 7VKT, 7WIH, 7WI8, 7WI6, 7VLA, 7VL9, 7VL8, 7TZF, 7TYY, 7TYW, 7TYO, 7TYL, 7TYH, 7TYF, 7T6V, 7T6U, 7T6S, 7T6T, 7TYX, 7TYN, 7TYI, 7VBH, 7EJX, 7RGP, 7RG9, 7RBT, 7RA3, 7WVY, 7WVX, 7WVW, 7WVV, 7WVU, 7EJK, 7EJA, 7EJ8, 7EJ0, 7EZC, 7SCG, 7SBF, 7W55, 7W57, 7W56, 7W53, 7WQ4, 7WQ3, 7SF7, 7WUJ, 7WUI, 7WUQ, 7SF8, 7WU4, 7WU3, 7WU5, 7WU2, 7U2L, 7XJH, 7XJI, 7EQ1, 7EPT, 7V68, 7V69, 7V6A, 7VQX, 7WBJ, 7X9C, 7X9B, 7X9A, 7SQO, 7EUO, 7WIG, 7WIC, 7SR8, 7S15, 7WKD, 7WV9, 7FEE, 7UL5, 7UL4, 7UL2, 7UL3, 7X8S, 7X8R, 7XTQ, 7RAN, 7WJ5, 7XBD, 7XBX, 7XBW, 7Y15, 7Y14, 7Y13, 7Y12, 7XOX, 7XOW, 7XOV, 7XOU, 7UM7, 7UM6, 7UM4, 7UM5, 7W0P, 7W0O, 7W0N, 7W0M, 7W0L, 7SUS, 8DCS, 8DCR, 7SK8, 7SK9, 7SK7, 7SK6, 7SK5, 7SK4, 7SK3, 7XTC, 7XTB, 7XTA, 7XT8, 7XT9, 7XMT, 7XMS, 7XNA, 7XN9, 7XMR, 7YXA, 7QVM, 7XKF, 7XKE, 7XKD, 8DU3, 7XW7, 7XW6, 7WU9, 7XW5, 7XZ6, 7XZ5, 7X9Y, 7XA3, 7YP7, 8DPI, 8DPH, 8DPG, 8DPF, 7X1U, 7X1T, 7XAV, 7XAU, 7XAT, 8CU7, 8CU6, 7R0C, 7X5H, 7VFX, 7SRS, 7SRR, 7SRQ, 7VIH, 7VIG, 7VIF, 7VIE, 7X2V, 7WZ7, 7WY0, 7WXW, 7WXU, 7W7E, 7W6P, 7YU8, 7YU7, 7YU6, 7YU5, 7YU4, 7YU3, 7XP6, 7XP5, 7XP4, 7Y89, 7F61, 7WZ4, 7WYB, 7WY8, 7WY5, 7PP1, 8DWH, 8DWG, 8DWC, 8EFQ, 8EFO, 8EFL, 8EFB, 8EF6, 8EF5, 7TS0, 7TRY, 7X10, 8E3Z, 8E3Y, 8E3X, 7XRR, 8E9X, 8EA0, 8E9Z, 8E9Y, 8E9W, 7U2K, 7T2H, 7T2G, 8F7X, 8F7R, 8F7Q, 8F7W, 8F7S, 8HAO, 8HA0, 8HAF, 7WCN, 7WCM, 7XW9 | 7B6W | |

**Table S2. Maxima of the apparent FRET distributions.**
Mean±SD for three technical replicas with different protein aliquots are given for $A_{2A}AR$ expressed in LEXSY or Sf9 and measured in apo form, in complex with antagonist ZM241385, or in complex with agonist NECA.

| | app. FRET | |
|---|---|---|
| | LEXSY | Sf9 |
| Apo | 0.60±0.01 | 0.57±0.01 |
| ZM241385 | 0.59±0.01 | 0.59±0.01 |
| NECA | 0.71±0.02 | 0.71±0.01 |

**Table S3. Fitting parameters for fFCS cross-correlation curves.**
fFCS cross-correlation curves were fitted with one anticorrelation term for the apo $A_{2A}AR$ and two anticorrelation terms for agonist-bound $A_{2A}AR$. $\tau_{diff}$ and $\tau_2$ were optimized globally across all four datasets. The half-widths of the 95% confidence intervals are given as fitting errors. $\chi_{red}^2 = 0.8$.

| | $\tau_{diff}$, ms | $\tau_1$, μs | $\tau_2$, ms | $A_1$ | $A_2$ |
|---|---|---|---|---|---|
| Apo LEXSY | 1.49±0.08 | 26±9 | 0.26±0.05 | 0.13±0.02 | - |
| NECA LEXSY | | 11±2 | | 0.55±0.07 | 0.35±0.03 |
| Apo Sf9 | | 41±17 | | 0.20±0.03 | - |
| NECA Sf9 | | 10±4 | | 0.30±0.08 | 0.40±0.04 |
